# YAP1 status defines two intrinsic subtypes of LCNEC with distinct molecular features and therapeutic vulnerabilities

**DOI:** 10.1101/2023.12.19.572449

**Authors:** C. Allison Stewart, Lixia Diao, Yuanxin Xi, Runsheng Wang, Kavya Ramkumar, Alejandra G. Serrano, B. Leticia Rodriguez, Benjamin B. Morris, Li Shen, Bingnan Zhang, Yan Yang, Samera H. Hamad, Robert J. Cardnell, Alberto Duarte, Moushumi Sahu, Veronica Y. Novegil, Bernard E. Weissman, Michael Frumovitz, Neda Kalhor, Luisa Solis Soto, Pedro da Rocha, Natalie Vokes, Don L. Gibbons, Jing Wang, John V. Heymach, Bonnie Glisson, Lauren Averett Byers, Carl M. Gay

**Affiliations:** Department of Thoracic/Head & Neck Medical Oncology, University of Texas MD Anderson Cancer Center, Houston, TX, USA; Department of Bioinformatics and Computational Biology, University of Texas MD Anderson Cancer Center, Houston, TX, USA; Department of Translational Molecular Pathology, University of Texas MD Anderson Cancer Center, Houston, TX USA; Lineberger Comprehensive Cancer Center, University of North Carolina at Chapel Hill School of Medicine, Chapel Hill, NC, USA; Department of Gynecologic Oncology and Reproductive Medicine, The University of Texas MD Anderson Cancer Center, Houston, TX, USA; Department of Medical Oncology, Hospital del Mar, Barcelona, Spain

**Keywords:** YAP1, LCNEC, EMT, AXL, DLL3, CD56

## Abstract

Large cell neuroendocrine carcinoma (LCNEC) is a high-grade neuroendocrine malignancy that, like the more common small cell lung cancer (SCLC), is associated with an absence of druggable oncogenic driver mutations, a clinically aggressive disease course, and dismal prognosis. In contrast to SCLC, however, there is little evidence to guide optimal treatment strategies which are, instead, often adapted from SCLC and non-small cell lung cancer (NSCLC) approaches. While there have been some efforts to describe the molecular landscape of LCNEC, to date there are few links between distinct biologic phenotypes of LCNEC and therapeutic vulnerabilities. Here, we demonstrate that the presence or absence of the transcription factor YAP1 distinguishes two roughly equal subsets of LCNEC. The YAP1-high subset is mesenchymal and inflamed and characterized, alongside *TP53* mutations, by co-occurring alterations in *CDKN2A/B* and *SMARCA4*. Therapeutically, the YAP1-high subset demonstrates vulnerability to MEK and AXL targeting strategies, including a novel preclinical AXL CAR-T cell, as well as predicted vulnerability to SMARCA2 degraders and CDK4/6 inhibitors. Meanwhile, the YAP1-low subset is epithelial and immune-cold and more commonly features *TP53* and *RB1* co-mutations, similar to those observed in pure SCLC. Notably, the YAP1-low subset is also characterized by expression of SCLC subtype-defining transcription factors -especially ASCL1 and NEUROD1 - and, as expected given its transcriptional similarities to SCLC, exhibits putative vulnerabilities reminiscent of SCLC, including Delta-like ligand 3 (DLL3) and CD56 targeting, as with novel preclinical DLL3 and CD56 CAR T-cells, and DNA damage repair (DDR) inhibition. These findings highlight the potential for YAP1 to guide the first personalized treatment strategies for LCNEC.

## Introduction

Neuroendocrine carcinomas (NECs) are clinically aggressive malignancies most commonly arising from the respiratory and gastrointestinal tracts, but, more rarely, from sites such as cervix, esophagus, urinary bladder, prostate, and ovary^1^. Their histologic appearance often resembles the small round blue cells of small cell lung cancer (SCLC), and, thus, are commonly classified as small cell carcinomas, while a minority of the cells are larger and classified as large-cell NECs (LCNECs) – together referred to as high-grade NECs (hgNECs). Pulmonary hgNECs account for approximately 20% of lung cancers and include SCLC, LCNEC, and combined histology NECs that may be difficult to distinguish histologically, particularly in small specimens collected via fine needle aspiration^2,3^. Inextricably linked histologically, these tumors are, thus, often managed via similar treatment paradigms. As SCLC is the most common of the pulmonary NECs, accounting for approximately 15% of all lung cancers compared to pulmonary LCNEC’s 1-3%^4^, the prevailing treatment paradigms are often derived from randomized trials including only SCLC patients^5^. In fact, these trials often actively exclude patients with LCNEC and, in many instances, combined histology NECs, as do many of the trials enrolling patients with non-small cell lung cancer (NSCLC). Owing to the lack of randomized clinical studies that permit pulmonary LCNEC or combined NEC histology, let alone those solely enrolling these tumors, few consensus clinical recommendations are available. Instead, clinical management varies according to treating physician and resemble standard of care approaches for SCLC or, less commonly, squamous NSCLC.

Typical pulmonary hgNEC patients are older with a history of heavy cigarette use^6^. These patients commonly present with distant metastases at diagnosis and are not amenable to definitive therapies such as surgery or radiation^4,7^. Due to a combination of demographics, comorbidities, and, especially, aggressive disease natural history, pulmonary hgNECs share a dismal prognosis with 5-year overall survival estimates ranging from just 5% (SCLC) to 15% (pulmonary LCNEC) across all stages^8^. While NSCLC patient survival has demonstrated a steady improvement with the continued development of personalized immunotherapies and targeted therapies, the prognoses for pulmonary hgNECs remain largely unimpacted by these developments^9^. Instead, metastatic pulmonary hgNEC patients are typically treated with a one-size-fits-all platinum-based chemotherapy approaches that yields robust initial responses, but are quickly undone by nearly inevitable resistance^10,11^. In metastatic SCLC, only modest benefit is observed with addition of immunotherapy and there are currently no predictive genomic biomarkers and zero approved targeted therapies^12,13^. The situation is similar in other hgNECs, aside from rare actionable oncogenic mutations (e.g. *KRAS* G12C) occasionally observed in pulmonary LCNEC^14^.

Recent evidence from SCLC suggests that the key to personalizing therapy for hgNECs may not be genomic biomarkers, despite their success in NSCLC. Instead, subtyping tumors on the basis of their transcriptome (or, similarly, their proteome, or their DNA methylome) provides a framework to delineate inter-tumoral heterogeneity. For example, in SCLC, transcriptional analyses reveal four subtypes – three defined by predominance of the transcription factors ASCL1 (SCLC-A), NEUROD1 (SCLC-N), and POU2F3 (SCLC-P), along with a fourth defined by inflammatory features (SCLC-I) such as antigen presenting machinery, T-cell infiltrate, and immune checkpoints^15^. These subtypes differ not only in their tumor immune microenvironment (TIME), but other spectra including EMT and neuroendocrine/non-neuroendocrine features. As for the potential clinical relevance of these subtypes, retrospective analyses demonstrate that SCLC-I may predict those patients most likely to experience long-term benefit from immunotherapy^15-17^, while preclinical data point to potential unique vulnerabilities among the other subtypes that might otherwise evade detection in unselected populations.

One issue is whether SCLC-derived transcriptional subtypes can be simply lifted wholesale from SCLC literature and applied to other hgNECs – both pulmonary and extrapulmonary. There is a scarcity of molecular data from other hgNECs and preliminary data in limited available cohorts that offer both support and opposition to this approach. The broadest effort thus far in pulmonary hgNECs excluding SCLC involved extensive molecular profiling of resected pulmonary LCNECs^18^. Though this profiling predated the current standard SCLC subtyping approach, heterogeneity among several features essential to SCLC subtyping (ASCL1 expression, neuroendocrine/non-neuroendocrine identity, inflamed/non-inflamed TIME) led to an initial subclassification of LCNEC into type I and type II categories. Like SCLC-A, type I LCNEC features high ASCL1 and neuroendocrine features, but an immune desert phenotype. Meanwhile, type II LCNEC possesses the low neuroendocrine, highly inflamed features more common to SCLC-I^18^.

In this report, we describe extensive molecular bulk and single-cell profiling of pulmonary LCNECs and combined histology NECs across multiple stages and with varied treatment histories in an effort to not only develop a classification schema for LCNECs but, more importantly, uncover important commonalities and distinctions across these malignancies with clinical relevance. Despite many shared molecular features with SCLC, treatment-naive pulmonary LCNECs and combined NECs warrant their own classification due to the unexpected prevalence of YAP1 as a defining feature in approximately 50% of cases. These YAP1-high NECs are defined not only by this transcriptional co-activator, but also relatively more mesenchymal and inflamed features, as well as unique genomic features, including mutually exclusive, frequent inactivation of *SMARCA4* and *CDKN2A/2B*. YAP1-high and -low hgNECs each have distinct therapeutic vulnerabilities owing to their unique genomic and transcriptomic features. Together, these findings demonstrate that YAP1 expression is a distinguishing feature of pulmonary LCNEC and combined histology with implications for inherent molecular and therapeutic features.

## Results

### YAP1 expression in pulmonary hgNECs

In order to evaluate YAP1 in pulmonary neuroendocrine tumors, including SCLC (n=18) and LCNEC (n=78) patients, we performed immunohistochemistry (IHC) using Clinical Laboratory Improvement Amendments (CLIA)-certified YAP1, ASCL1, NEUROD1, and POU2F3 assays. Mixed histology tumors included combinations of LCNEC with SCLC, adenocarcinoma, and squamous cell carcinoma. Scoring of IHC expression was performed in malignant cells and specifically in the NEC cells of the mixed histology tumors. Abundant nuclear YAP1 expression was detected in approximately half of the LCNEC tumors and a subset of the mixed histology tumors and these frequently occurred in the absence of ASCL1, NEUROD1, or POU2F3 (Figure 1a-b). There was no difference in nuclear YAP1 H-score values between treatment-naïve (naïve) and previously treated (treated) LCNEC patients (Figure 1c). There were not enough samples from relapsed SCLC or mixed histology patients in our cohort to perform a similar analysis. In fact, some of the highest YAP1 H-score values were detectable in treatment-naïve LCNEC tumors (Figure 1d). The largely mutual exclusive levels of ASCL1 and YAP1 were confirmed in LCNEC xenograft models with similar histology (Figure 1e). YAP1 protein, includingH-score values and reverse phase protein array (RPPA) levels and mRNA levels were highly concordant across LCNEC cell lines and patient tumors and bimodally distributed, based on bimodal index^19^ (Extended data figure 1b). The bimodal index was used to determine YAP1-high and -low LCNEC subsets. Additionally, YAP1, phosphorylated S127 YAP1, and TAZ levels by reverse phase protein array (RPPA) were higher in YAP1-high compared to YAP1-low LCNEC cells, as determined by bimodal index (Extended data figure 1c).

**Figure 1.**
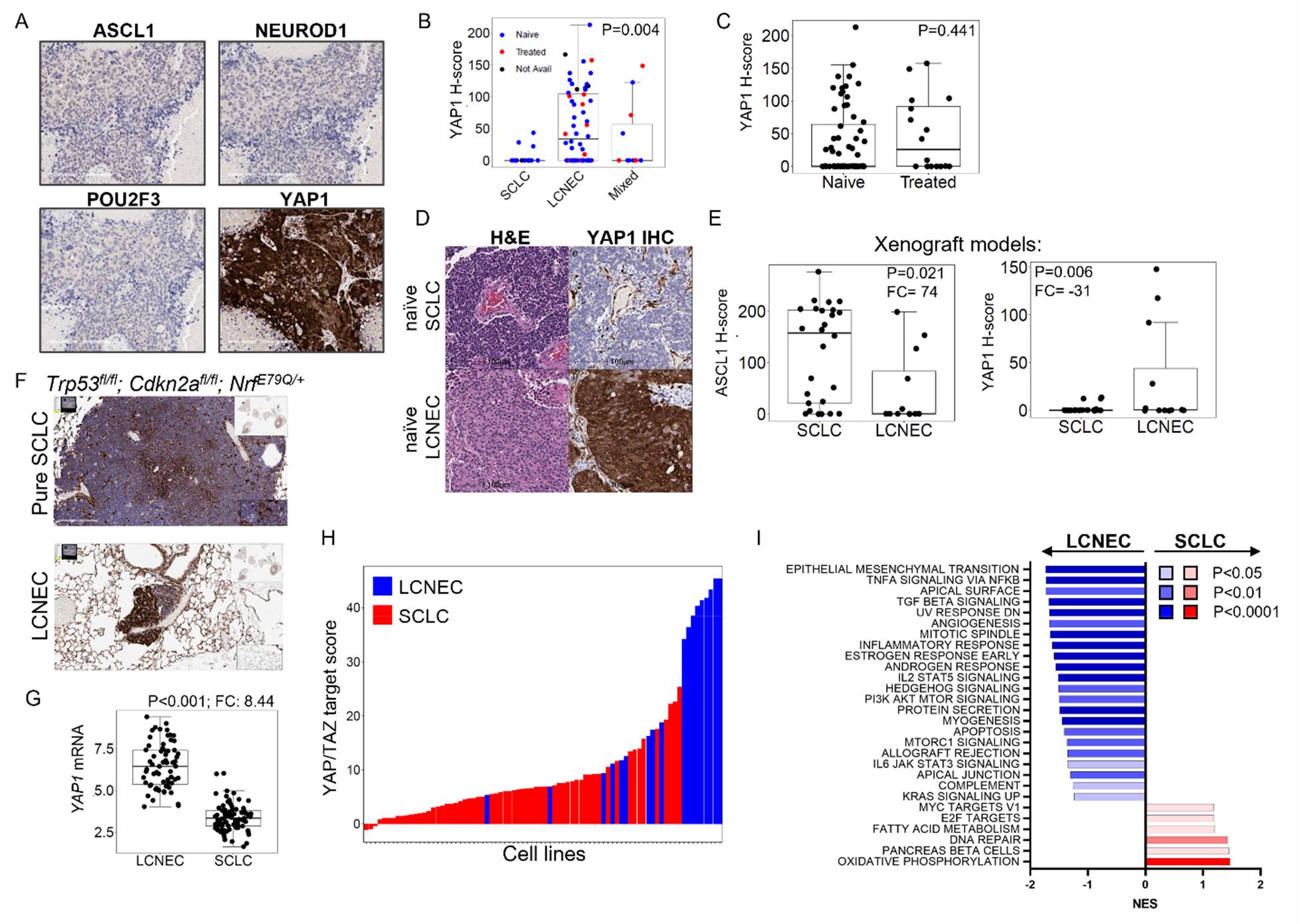
YAP1 is abundant in LCNEC, regardless of treatment status. A, Representative IHC for ASCL1, NEUROD1, POU2F3, and YAP1 in a LCNEC surgical resection. B, YAP1 nuclear H-score values in patient tumors colored by treatment status (naïve, treated, not available). C, YAP1 nuclear H-score values are not different in naïve and treated LCNEC tumors. D. Representative IHC for YAP1 in treatment naïve SCLC and LCNEC. E, ASCL1 H-scores are higher in SCLC PDX models and YAP1 H-scores are higher in LCNEC PDX models. F, Representative IHC demonstrating nuclear YAP1 in the LCNEC cells, but not SCLC cells from GEMMs with combined histology tumors. Insets demonstrate lower magnification images. G, *YAP1* expression is higher in LCNEC compared to SCLC tumors. I, YAP/TAZ target score is higher in LCNEC cell lines compared to SCLC. J, GSEA of LCNEC and SCLC patient tumors. Scale bar = 100 μm.

### LCNEC specificity of YAP1

In order to determine if nuclear YAP1 is LCNEC specific, YAP IHC was performed using a unique *Trp53*^*f/f*^; *Cdkn2a*^*f/f*^; *Nrf*^*E79Q/+*^ genetically engineered mouse model with SCLC/LCNEC combined histology^20^. Nuclear YAP1 was only detectable in LCNEC cells (Figure 1f). *YAP1* mRNA levels are higher in LCNEC compared to SCLC patient tumors (Figure 1g). LCNEC cell lines have higher levels of YAP1 and phosphorylated S127 YAP1 compared to SCLC cell lines (Extended data figure 1d). A YAP/TAZ transcriptional target signature of 22 genes that robustly infers pathway activity^21^ was applied to LCNEC and SCLC cell lines and the highest activity was detectable in LCNEC cell lines (Figure 1h). Gene set enrichment analysis (GSEA) of LCNEC and SCLC patient tumors revealed an enrichment in EMT, TNF-α and TGF-β signaling in LCNEC tumors and oxidative phosphorylation and DNA repair in SCLC tumors (Figure 1i). Notably, this is consistent with previous data demonstrating an enrichment in DNA damage repair signaling in SCLC^22^. These data demonstrate that YAP1 levels and activity are enriched in LCNEC compared to SCLC.

### Distinct genomic alterations by YAP1 status

In order to determine genomic differences between YAP1-high and -low LCNEC, we evaluated common alterations in whole genome sequencing of 21 LCNEC cell lines in the Cancer Cell Line Encyclopedia^23^. YAP1-high cell lines commonly have *CDKN2A/B* homozygous deletions (67%/47%) and *SMARCA4* mutations (Figure 2a; 40%). These alterations were not detectable in YAP1-low cell lines. While *RB1* mutations were present in both YAP1-high and -low cell lines, intact *RB1* was more common in the YAP1-high cells (87% vs. 50%). There was no difference in frequency of *TP53* mutations across cell lines. Consistent with these genomic alterations, protein levels of p16, protein encoded by *CDKN2A*, were lower in LCNEC cell lines compared to SCLC cell lines. Conversely, RB1 protein levels were higher in LCNEC compared to SCLC cell lines (Figure 2b), but within LCNEC, RB1 is higher in YAP1-high cell lines and patient tumors (Figure 2c). *RB1* was positively correlated with *YAP1* (Figure 2d). Even a minor knockdown of RB1 directly reduced YAP1 levels (Figure 2e). In contrast, expression of *CDKN2A* and *SMARCA4* was negatively correlated with *YAP1* in LCNEC patient tumors (Figure 2f). Interestingly, despite the frequent inactivating mutations of *SMARCA4* in YAP1-high LCNEC, its expression was higher overall in LCNEC compared to SCLC (Extended data figure 1f). This is likely because *SMARCA4* mutations create a dominant negative form of the protein, as most of the mutations are truncations, and this is consistent with inactivating hotspot mutations in *TP53*^24^.

**Figure 2.**
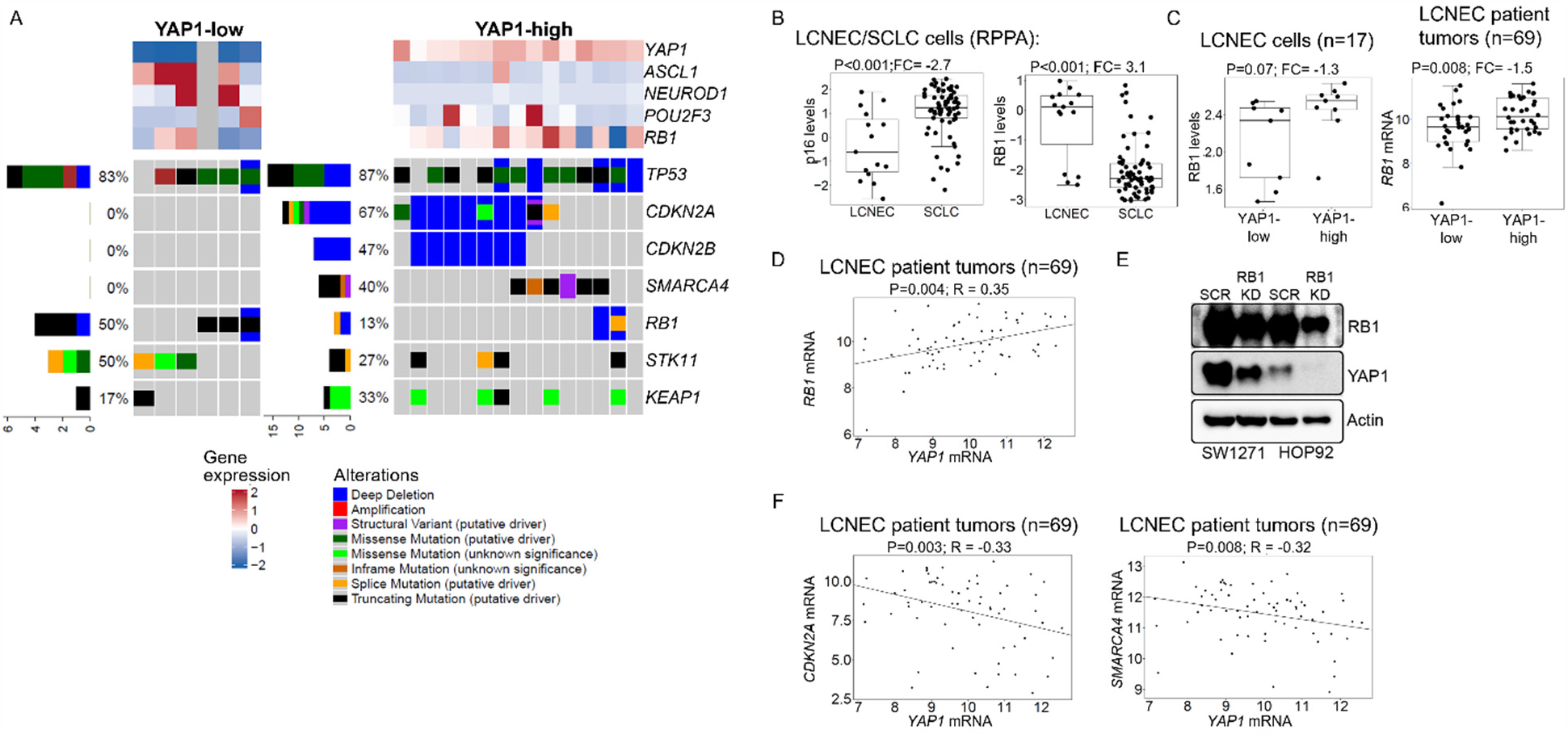
YAP1-high LCNEC is genomically distinct from YAP1-low LCNEC. A, Genomic alterations in YAP1-high and low LCNEC cell lines, including frequency of mutations (left), gene expression of YAP1, RB1, and SCLC molecular subtype markers. B, RPPA protein levels of p16 and RB1 in SCLC and LCNEC cell lines. C, RPPA protein levels (left) and mRNA (right) of RB1 in YAP1-high and -low LCNEC cell lines (left) and patient tumors (right). D, *RB1* expression is correlated with *YAP1* in LCNEC patient tumors. E, When RB1 levels are knocked down, YAP1 is also reduced. F, *CDKN2A* and *SMARCA4* are negatively correlated with *YAP1* expression in LCNEC patient tumors.

### Features of YAP1-high LCNEC

To gain a greater understanding of YAP1 subsets in LCNEC patient and cell line cohorts, transcriptional differences were analyzed. Similar to protein levels, *YAP1* and *ASCL1, NEUROD1*, and *POU2F3* were largely mutually exclusive (Figure 3a). As expected, the YAP/TAZ transcriptional target score was elevated in the YAP1-high samples (Figure 3a), as well as other YAP1-associated genes (Extended data figure 2a-c). Additionally, the lung EMT score^25^ was elevated in the YAP1-high samples (Figure 3a; Extended data figure 2c), indicating that these samples were more mesenchymal. Along these same lines, the neuroendocrine (NE) score was reduced in the YAP1-high samples (Figure 3a; Extended data figure 2c). This is consistent with data in SCLC, where EMT score and NE status are inversely related^15^. In fact, siRNA knockdown of YAP1 resulted in a similar loss of vimentin levels and an increase of synaptophysin, a common neuroendocrine marker, suggesting that YAP1 plays a direct role in the neuroendocrine and EMT status of LCNEC cells (Figure 3b). Cancer Cell Line Encyclopedia reduced representation bisulfite sequencing (RRBS)^23^ of LCNEC cell lines revealed that YAP1 is methylated in YAP1-low cell lines and ASCL1 is methylated in YAP1-high cell lines, indicating that expression of these subtype-defining factors is modulated epigenetically (Extended data figure 2d). RPPA analysis of LCNEC cell lines demonstrated higher expression of YAP1, phospho-YAP1, NOTCH2, and vimentin in the YAP1-high cell lines and ASCL1, DLL3, E-cadherin, gH2AX and BCL2 in the YAP1-low cell lines (Figure 3d). Consistent with this, EMT, inflammatory pathways, TNF alpha, TGFB and NOTCH signaling are enriched in the YAP1-high patient tumors and DNA repair, E2F targets and oxidative phosphorylation are abundant in the YAP1-low tumors.

**Figure 3.**
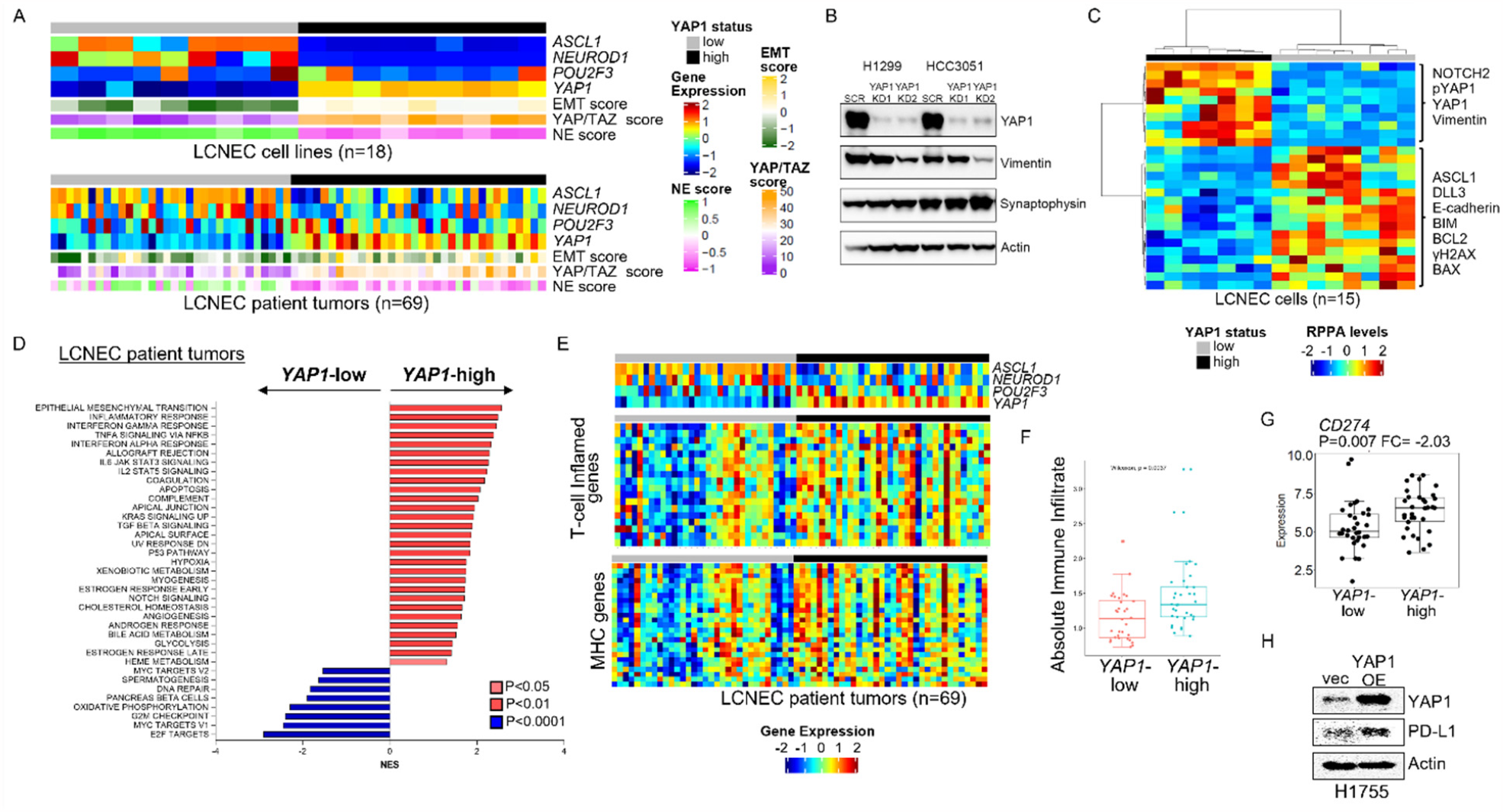
YAP1 defines distinct subsets of LCNEC. A, YAP1-high LCNEC cells (top) and tumors (bottom) have a higher YAP/TAZ target score and EMT score, but lower NE score. B, YAP1 knock down results in a decrease in vimentin and increase in synaptophysin levels. C, Heatmap of RPPA protein differences in YAP1-high and low cells. D, GSEA of YAP1-high and - low patient tumors. E, Heatmap of T-cell inflamed and MHC genes based sorted by YAP1-status. F. CIBERSORT analysis of YAP1-high and -low patient tumors. G, YAP1-high LCNEC patient tumors have higher expression of *CD274* (PD-L1). H, Similarly, when YAP1 was overexpressed, PD-L1 levels are increased.

In order to further investigate inflammation in LCNEC, expression of genes included in the Tumor Inflammation Score (TIS)^26^ were evaluated in YAP1-high and -low patient tumors (Figure 3e). YAP1-high LCNEC is reminiscent of SCLC-I tumors, with abundant expression of a T-cell inflamed signature and MHC genes, while there is also a subset of the YAP1-low tumors with an abundant inflammatory signature. Specifically, YAP1-high tumors have increased absolute immune infiltrate by CIBERSORTx compared to YAP1-low (Figure 3f). Consistent with increased inflammation, YAP1-high tumors had higher expression of PD-L1 (*CD274*). When YAP1 was overexpressed in a YAP1 low LCNEC cell line, PD-L1 levels were similarly increased (Figure 3g,h) suggesting that YAP1-status may predict response to immune checkpoint blockade in LCNEC. YAP1-high tumors additionally have high expression of *MICA* and NKG2A (*KLRC1)* (Extended data figure 2e), suggesting that these tumors may be more responsive to natural killer (NK) cell killing and/or monalizumab targeting (anti-NKG2A).

### Single-cell transcriptional profiling of LCNEC patient samples and PDX models

In order to further understand the heterogeneity of *YAP1* expression and activity in relapsed LCNEC patient biopsies and established PDX models, single-cell RNAseq was performed. Six core needle biopsies and surgical resection from patients with pLCNEC or combined histology tumors, including LCNEC, were dissociated and analyzed by single-cell RNAseq. A total of 18,378 cells passed the initial QC analysis and were classified by cell type (Figure 4a). The 8,069 pooled cancer cells were selected and color-coded by patient (Figure 4a). Similarly, single-cell RNAseq was performed on 15,246 cells from six PDX tumors from models derived from patients diagnosed with pLCNEC or combined histology tumors, including LCNEC (Figure 4b). In contrast to previous data with SCLC PDX models^27^, cells from different LCNEC patient biopsies were intermixed, which indicates that LCNEC tumors and PDXs have similar biologic phenotypes. To determine expression of common genes in individual patient or PDX sample cancer cells, binary expression of *YAP1, ASCL1, UCHL1* (NE marker) and *REST* (non-NE marker) were determined and visualized on Uniform Manifold Approximation and Projection (UMAP) plots (Figure 4d; Extended data figure 3a). *YAP1* was expressed by more than 15% of cells in three or more patient biopsies or PDX tumors (Figure 4c), which mirrors the YAP1 IHC data in LCNEC tumors. Similar to bulk protein and mRNA, samples with a higher percentage of *YAP1* positive cells also have higher *REST* and lower *ASCL1* and *UCHL1* and vice versa (Extended data figure 3a). *ASCL1* and *YAP1* were not frequently co-expressed by cells. Consistent with bulk RNAseq data, *YAP1*-positive cells express abundant *REST* compared to *YAP1*-negative cells. While YAP1 is associated with lower NE gene expression, *UCHL1* was expressed by both *YAP1*-positive and -negative cells which is further validation that these are truly NE cells (Extended data 3b). Additionally, binary gene expression and EMT score for pooled cancer cells were visualized on UMAP plots and demonstrate a high EMT score in *YAP1*-positive cell populations (Figure 4d). As expected, similar expression was found in a patient biopsy (MDA-SC288A) and PDX derived from the same biopsy (MDA-SC288APDX) (Extended data figure 3c). Furthermore, single-cell RNAseq from the MDA-SC271-2 circulating tumor cell derived xenograft (CDX) model mirrors YAP1 and cMYC IHC protein levels for the same model (Extended data figure 3d). Consistent with GSEA analysis of YAP1-high and -low LCNEC patient tumors, single-cell analyses revealed an enrichment of inflammatory response pathways, TNF-α and TGF-β, and EMT in the YAP1-positive cell population from LCNEC patient biopsies (Figure 4e).

**Figure 4.**
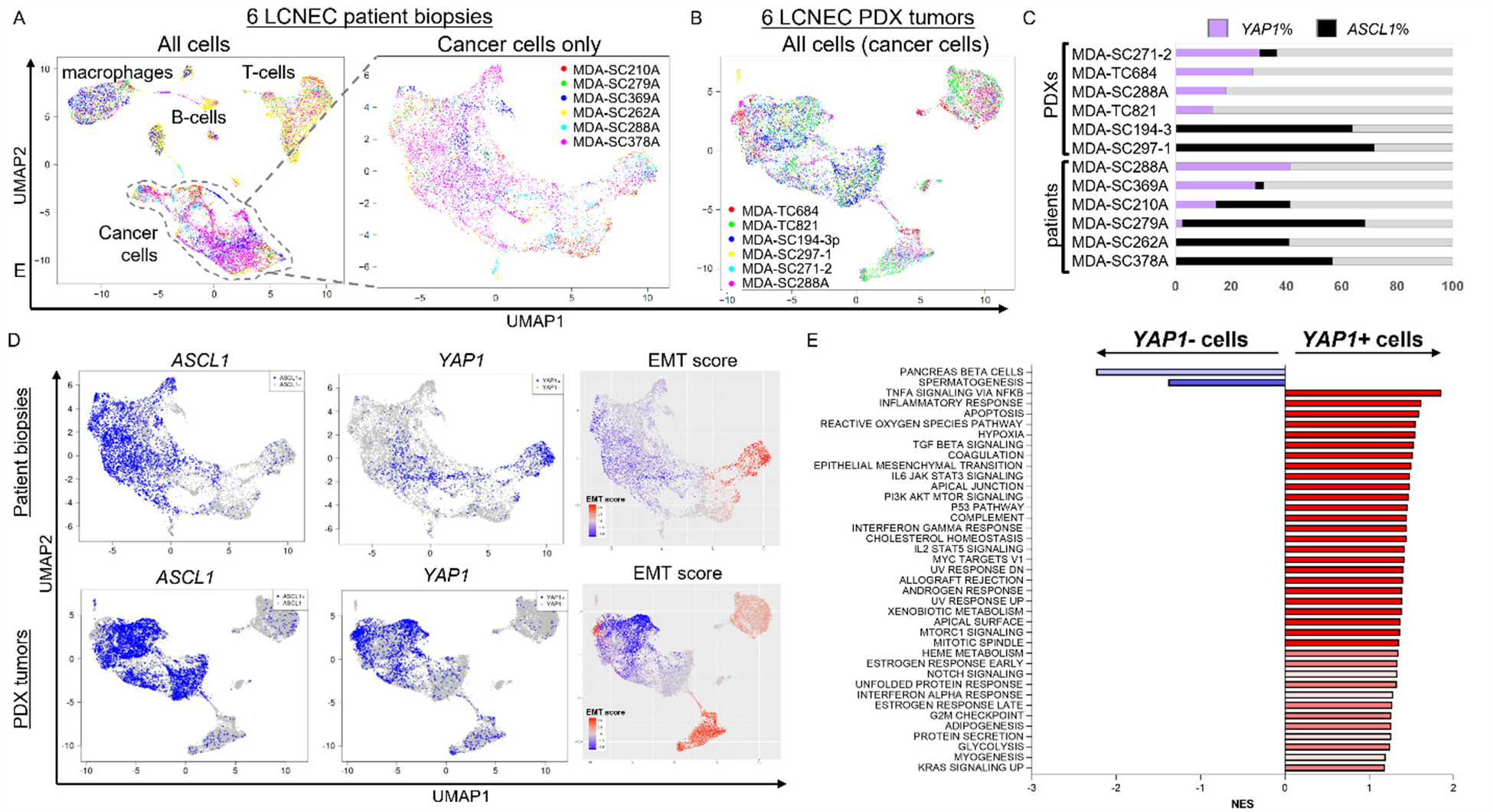
YAP1 in LCNEC patient biopsies and PDX models is associated with higher EMT. A, UMAP plot of all cell populations from 6 LCNEC and combined histology biopsies (left) and UMAP plot of cancer cell populations (right), both labeled by patient. B, UMAP plot of all cell populations from 6 LCNEC PDX models, labeled by model. C, Percentage of YAP1-positive cells in each biopsy. D, UMAP plots demonstrating binary expression of ASCL1 and YAP1. YAP1-expression cell populations had elevated EMT scores. E, GSEA of *YAP1*-positive and - negative cells.

### Therapeutic response in LCNEC preclinical models

Platinum chemotherapy remains the backbone of treatment for LCNEC – regardless of whether SCLC or NSCLC paradigms are followed. Response to platinum chemotherapy was tested in preclinical models, including LCNEC cell lines and PDX models (Figure 5a,b; Extended data figure 4a), and YAP1 status did not predict response to cisplatin. As for YAP1-specific approaches, there are multiple inhibitors of YAP1/TEAD association available (e.g., verteporfin, MYF-01-37, XAV-939, TED347). Additionally, verteporfin was able to reduce YAP1 levels (Figure 5c). However, overexpression of YAP1 did not alter verteporfin response (Extended data figure 4b) and verteporfin (Figure 5d) and TED347 (Extended data figure 4c) were most effective only at reducing cell proliferation and independent YAP1 status. For a broader investigation of YAP1-dependent LCNEC vulnerabilities, Cancer Cell Line Encyclopedia (CCLE) Cancer Dependency Portal (DepMap) data was analyzed for 15 YAP1-high and 6 YAP1-low LCNEC cell lines for drug response (Figure 5e). As a breakdown of drug classes, YAP1-high cell lines were more sensitive to trametinib, selumetinib, refametinib (MEK1/2 inhibitors), palbociclib (CDK4/6 inhibitor), dasatanib (SRC family kinase inhibitor), dabrafenib (BRAF inhibitor for *BRAF* V600E mutation), FTI-277 (farnesyl transferase inhibitor), JNK-IN-8 (JNK 1/2/3 inhibitor), tozasertib (AURKA inhibitor), vinblastine (inhibits microtubule assembly), bleomycin (antibiotic). In contrast, YAP1-low LCNEC cell lines were more sensitive to gemcitabine (antimetabolite chemotherapy), fulvestrant (estrogen receptor antagonist), docetaxol (taxane chemotherapy), temozolomide (alkylating chemotherapy), venetoclax (BCL2 inhibitor), tamoxifen (selective estrogen receptor modulator), entinostat (histone deacetylase 1/3 inhibitor), and gefitinib (EGFR inhibitor). Of these, YAP1-high cell lines were sensitive to multiple MEK1/2 inhibitors and trametinib was selected for additional analyses. Single-agent trametinib was unable to consistently alter YAP1 levels (Figure 5f). However, overexpression of YAP1 was able to enhance response to trametinib in a YAP1-low cell line (Extended data figure 4d). Following reports of synergistic activity of trametinib in other cancers, combinations with cisplatin and palbociclib were tested with no additive effect detected, regardless of YAP1 status (Extended data figure 4e). Trametinib single-agent delayed tumor growth in two YAP1-high, rapidly growing, cell line xenograft models. H1915 is a LCNEC cell line and SW1271 is a SCLC cell line with a YAP1-fusion, suggesting that MEK1/2 inhibition may be effective in high-grade neuroendocrine carcinomas with high YAP1 levels, regardless of small or large cell morphology (Figure 5g). However, there remains potential to improve response to trametinib using combination therapies.

**Figure 5.**
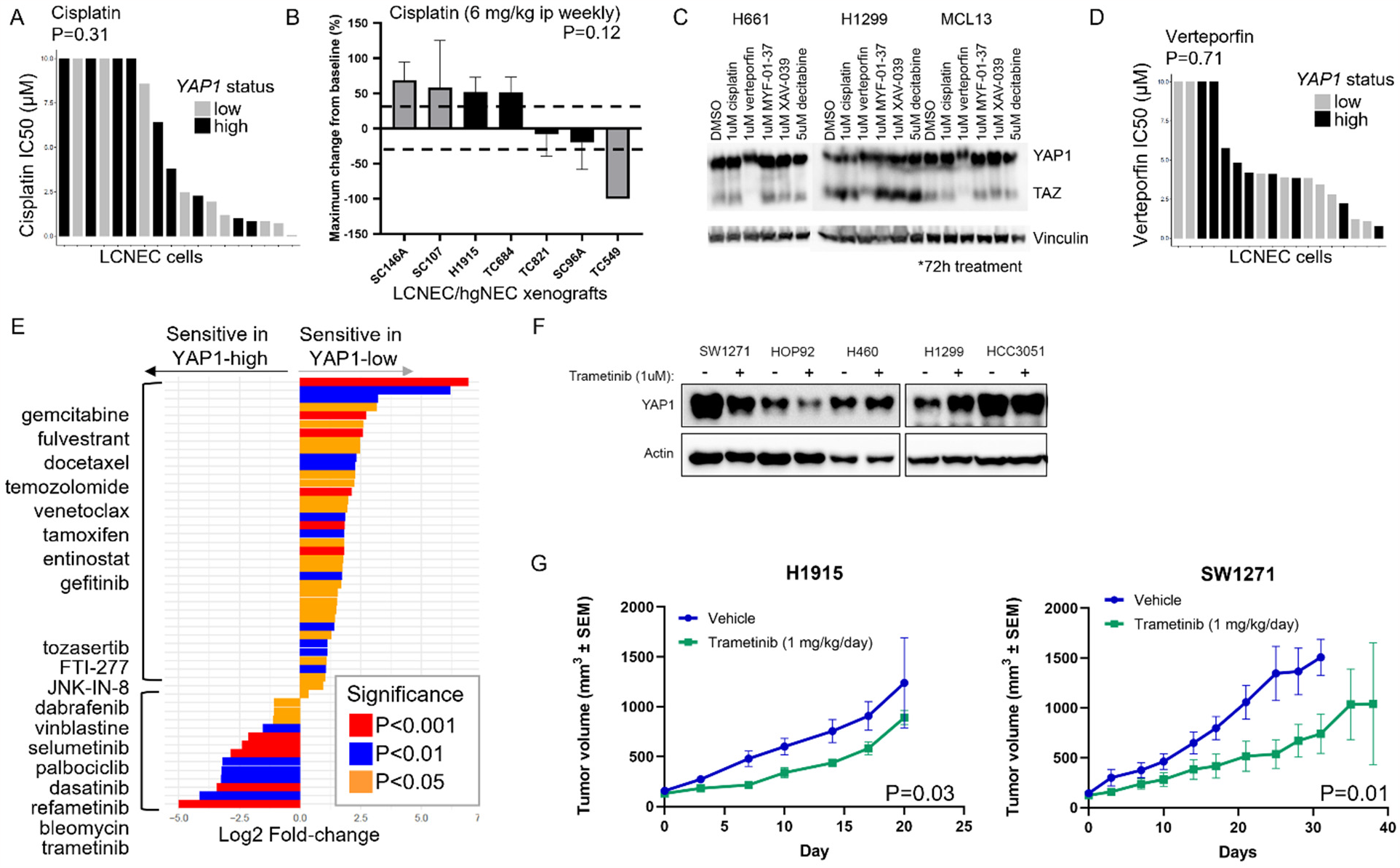
YAP1-high LCNEC is sensitive to MEK1/2 inhibition. YAP1 status does not predict sensitivity to cisplatin in LCNEC cell lines (A) or cell line/PDX xenograft models (B). C, Treatment of LCNEC cell lines with DMSO, cisplatin, MYF-01-37, XAV-939 and decitabine did not change YAP1, but verteporfin reduced YAP1 levels. D, YAP1 status does not predict response to verteporfin. E, Waterfall plot of drug sensitivity in LCNEC cell lines. A comparison of high YAP1 levels and IC50 values identified several drugs with similar targets, including MEK1/2, CDK4/6, and Src family kinase inhibitors. F, Treatment of YAP1-high LCNEC and SCLC (SW1271) cell lines with trametinib did not change YAP1 levels. G, Tumor growth curves from YAP1-high cell line xenografts H1915 (LCNEC) and SW1271 (SCLC) demonstrate a delay in tumor growth with trametinib treatment.

### Surfaceome Targeting

While there may exist differences in response to established anti-cancer therapies, the distinct features of YAP1-high and -low LCNEC could assist in identifying novel therapeutics. Several emerging anti-cancer strategies exploit the tumor cell surfaceome to deliver cytotoxics (antibody-drug conjugates [ADCs]) or immune cells (bi-specific T-cell engagers [BiTEs] or chimeric antigen receptors [CARs]). While none of these are currently approved for LCNEC/SCLC, several have been explored and shown activity--including a CAR-T (AMG119)^28^ and a BiTE (tarlatamab) targeting Delta-like ligand 3 (DLL3)^29,30^. Expression of known cell surface targets in other cancer types were evaluated in LCNEC tumors and *CD22, CD33*, HER2/*ERBB2*, TROP2/*TACSTD2* were elevated in YAP1-high tumors (Extended data figure 5a). Due to previous reports of common cell surface targets in SCLC^29-32^ or NSCLC^33^, alternative cell surface targets investigated in LCNEC include, DLL3, AXL, CD56, and EPCAM. Notch inhibitory ligand DLL3, is heterogeneously expressed across SCLC, enriched particularly in the non-inflamed subtypes. AXL is a transmembrane receptor tyrosine kinase known to promote EMT and therapeutic resistance in NSCLC^25,33^. CD56 (*NCAM1*) is a glycoprotein that is expressed on the surface of SCLC and other neuroendocrine tumors and commonly used for diagnosis by IHC. Epithelial cell adhesion molecule (EPCAM) is abundantly expressed by the cell surface of SCLC cells and is commonly used as a marker to identify or capture circulating tumor cells. AXL mRNA and protein are abundantly expressed by YAP1-high LCNEC patient tumors and cell lines (Figure 6a; Extended data figure 5b). In contrast, DLL3 and CD56/NCAM1 were more abundant in the YAP1-low LCNEC patient tumors and cell lines. *EPCAM* was highly expressed by LCNEC tumors, regardless of YAP1 status (Extended data figure 5c). LCNEC cell lines were analyzed by flow cytometry for cell surface expression of AXL, DLL3, and CD56. AXL was more abundant on the cell surface of YAP1-high LCNEC cell lines (Figure 6b). While there was no difference in cell surface DLL3, CD56, or EPCAM between YAP1-high and -low LCNEC cell lines, detectable levels were present in most cell lines evaluated (Figure 6b, Extended data figure 5d).

**Figure 6.**
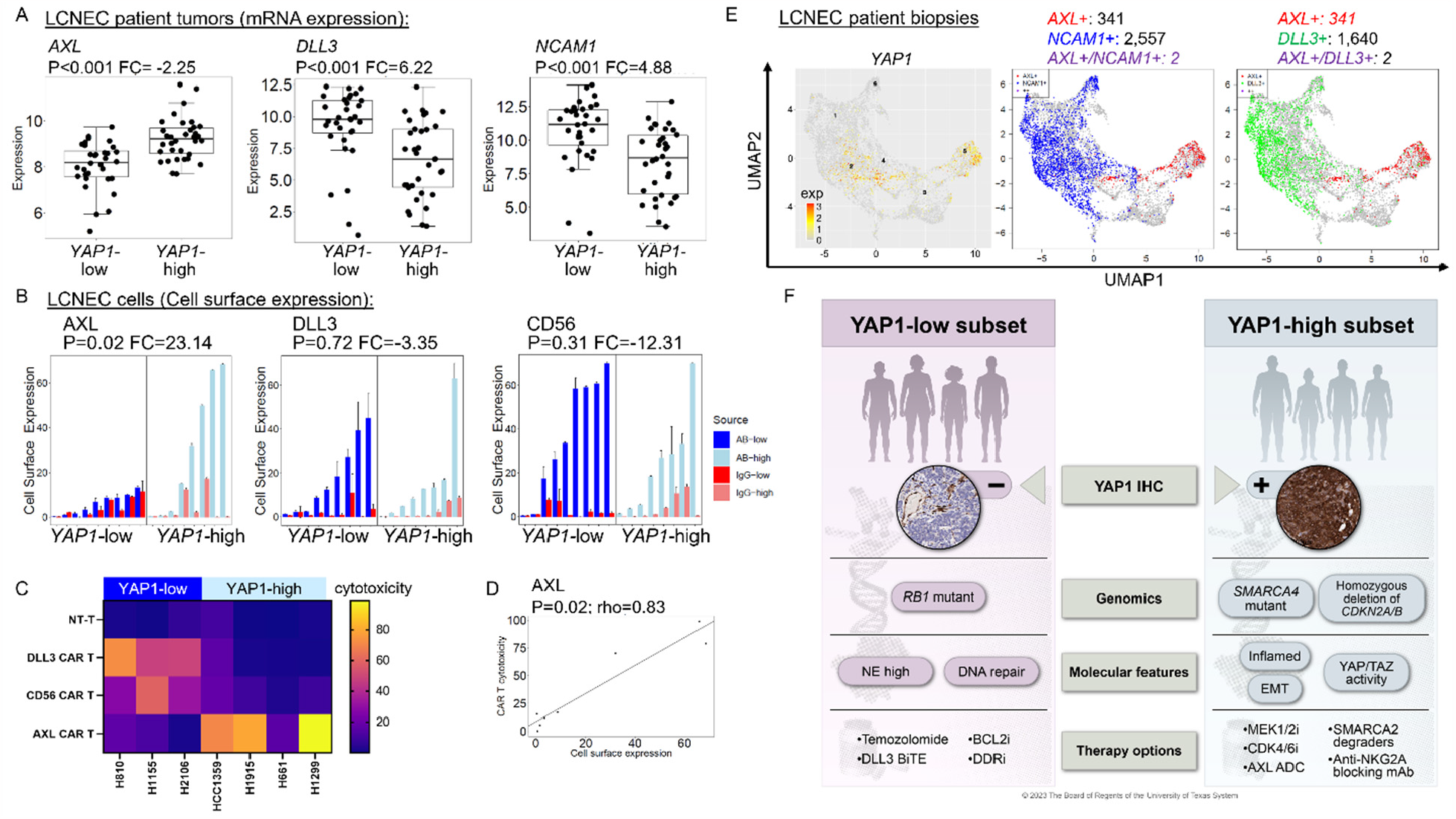
Cellular therapies targeting YAP1 subsets of LCNEC. A, *AXL* is expressed by YAP1-high and *DLL3/NCAM1* are expressed by YAP1-low patient tumors. B, Cell surface expression of AXL, DLL3 and CD56 in LCNEC cell lines by flow cytometry demonstrates a similar pattern. C, Cytotoxicity of NT-T, DLL3 CAR-T, CD56 CAR-T, and AXL CAR-Ts in subsets of LCNEC. D, Correlation between cell surface expression of AXL and cytotoxicity of AXL CAR-T in LCNEC cell lines. E, UMAP plots visualizing YAP1 (left) or AXL (red) and NCAM1 (blue) or AXL (red) and *DLL3* (green) in 6 LCNEC patient biopsies. Very few cells express both *DLL3* and *AXL* or *NCAM1* and *AXL*, suggesting that distinct cell populations can be targeted via the surfaceome. F, Schematic demonstrating common features of YAP1-high and low subtypes of pulmonary LCNEC.

To test the approach of targeting the cell surface in LCNEC, third-generation CARs against DLL3, CD56, and AXL were generated that consist of dual co-stimulatory domains (CD28 and 4-1BB); a target-specific single-chain variable fragment (scFv); IgG2 CH2 and CH3 domains with a N297Q mutation; and the CD3ζ intra-cellular domain. DLL3 and CD56 CAR-Ts were effective at inducing cytotoxicity in YAP1-low LCNEC cell lines, while AXL CAR-Ts were effective in YAP1-high LCNEC cell lines (Figure 6c). In LCNEC cell lines, AXL CAR-Ts were most effective in cell lines with high AXL (and YAP1) levels (Figure 6d; Extended data figure 5e), which is likely due to close interaction between AXL and YAP1. In fact, knockdown of YAP1 reduces AXL expression in LCNEC cell lines (Extended data figure 5f). While expression of cell surface targets is relatively discrete in cell lines, allowing for clear differences in response to cellular therapies (Figure 6b,c), the single cell data shows heterogeneity of expression. Specifically, data from the patient biopsies demonstrates surface target expression by largely mutually exclusive cellular populations (e.g., *AXL* and *DLL3* or AXL and *NCAM1*) even in the same biopsy (Figure 6e; Extended data figure 5g,h). However, this may suggest that pooling ADCs/BiTES/CAR-Ts targeting distinct cellular populations may be a novel way to improve response in recalcitrant pulmonary hgNECs. Overall, pulmonary LCNEC consists of two distinct, roughly equal, subtypes that can be identified with a CLIA-certified, nuclear YAP1 IHC assay, each with defining genomic and molecular features, and specific known and predicted therapeutic vulnerabilities (Figure 6f).

## Discussion

Pulmonary hgNECs have been plagued by classifications both too rigid, as with the frequent exclusion of LCNECs and combined histology patients from many trials, and too flexible, as with the tendency to treat all SCLC or even all hgNECs according to a single paradigm. In part, this reflects the challenge of collecting sufficient tissue from these tumors due to their rarity and even more rarely, the collection of core biopsies and surgical resections for research purposes. This paucity of samples is further amplified in the relapsed setting – a setting in which there has never been any standard indication for repeat biopsy following initial relapse. As a result, practitioners have been left with slowly evolving, all-comers approaches to disease entities with ever-clearer inter- and intra-tumoral heterogeneity.

YAP1 status clearly delineates two subsets of pulmonary LCNEC that feature distinct biology well beyond their YAP/TAZ pathway activation. The YAP1-high tumors exhibit low neuroendocrine features alongside high mesenchymal features and a relatively more inflamed TIME in contrast to the more epithelial, neuroendocrine, and immune-cold YAP1-low tumors. YAP1 status also demarcates those YAP1-low tumors with almost exclusively classic SCLC-like genomic features (i.e. dual loss of *TP53 and RB1*) and YAP1-high tumors with a variety of other genomic alterations less commonly observed in *pure* SCLC (e.g. *CDNK2A/B, SMARCA4*).

While delineating this disease into two subsets is biologically provocative, it is critical to consider the implications on current and, especially, future therapeutic approaches. While management of LCNEC often follows a mix of SCLC and NSCLC paradigms, there is no evidence to support any selection between these paradigms on the basis of any molecular feature aside from exceedingly rare instances of actionable driver mutations a la NSCLC. Furthermore, LCNEC patient outcomes are uniformly poor, seemingly regardless of the paradigm adopted for their treatment, which calls into question the use of *either* the SCLC or NSCLC paradigms as optimal. In other words, it is unlikely a matter of an elusive biomarker for selection between a platinum-etoposide or platinum-taxane doublet as the backbone for LCNEC patient chemo-immunotherapy. Instead, a whole new paradigm is needed and these data argue that investigators should consider LCNEC and similar mixed histology hgNECs distinctly on the basis of, first and foremost, the tumor’s YAP1 status.

These findings suggest that approved (e.g., PD-(L)1 or CTLA4) or investigational (NKG2A) targets for immune checkpoint blockade would be relatively more effective if only YAP1-high patients were selected. Similarly, consideration of cell surface molecules that could be exploited as beacons for delivery of cytotoxic payload (ADCs) or immune cells (CARs, BiTEs) should be considered on a YAP1-dependent basis, as demonstrated with CAR T-cells targeting DLL3, CD56, and AXL. Even the genomic distinctions between YAP1-high and -low uncover potential vulnerabilities, including SMARCA2 degraders (for *SMARCA4*-deficient tumors) or CKD4/6 inhibitors (for *CDKN2A/B*-deficient tumors) that will likely only expand with development of novel targeted therapies. LCNECs are rare but not so rare that it should preclude a community effort to review responses to standard-of-care chemo-immunotherapy with a single retrospective biomarker and certainly not so rare that prospective, biomarker-driven trials should not be pursued with current or future therapies.

## Methods

### Transcriptional sequencing datasets

RNAseq analysis datasets from normal tissues were obtained from GTEx Portal (http://www.gtexportal.org) and tumor samples were retrieved from published and unpublished datasets in multiple human tissues, including SCLC^34^, LCNEC^18^. Cell line data were obtained from multiple sources, including CCLE LCNEC cell lines (n=15)^35-37^ and SCLC (n=62)^38^.

### Cell culture

All cell lines were grown in RPMI with 10% fetal bovine serum and antibiotics and cultured at 37°C in a humidified chamber with 5% CO2. All cell lines were frequently tested for Mycoplasma. Cell lines demonstrated a range of phenotypic characteristics, from floating aggregates to spindle-like or cobblestone attached, but these were consistent with ATCC or previous reports.

### Patient consent and tissue collection

Patients diagnosed with SCLC, LCNEC, or a combined high-grade neuroendocrine carcinoma at the University of Texas M.D. Anderson Cancer Center were selected on the basis of disease irrespective of age, gender or other clinical criteria. These patients underwent informed consent to Institutional Review Board (IRB)-approved protocol 2020-0412 or PA14-0276. Core needle biopsies, EBUS biopsies, or surgical resections were collected into RPMI media with antibiotics and transported quickly to the research lab for sample preparation and single-cell RNAseq.

### Histology and IHC

Independent pulmonary pathologists carefully reviewed H&E images from LCNEC, SCLC, and combined histology tumors to accurately assess morphology. Immunohistochemistry was performed with a Bond Max automated staining system (Leica Microsystems Inc., Vista, CA) using a CLIA certified YAP1 antibody and protocol used by the MDACC Translational Molecular Pathology Department for clinical samples. Nuclear expression YAP1 was quantified using a 4-value intensity score (0, none; 1, weak; 2, moderate; and 3, strong) and the percentage (0%– 100%) of reactivity. A final expression score (H-score) was obtained by multiplying the intensity and reactivity extension values (range, 0–300), as described previously^22,39^. IHC data were examined by two experienced pathologists (LSS, JF).

### Flow Cytometry

One million cells each for H810, H1155, H1755, H1833, H2106, MKL1, HCC2374, HCC4017, HOP92, H1359, H2066, H1770, H2106, H1570, H661, HCC3051, H460, HCC4017, H1299, and H1915 in triplicate were surfaced stained with DLL3 (FAB4315P; R&D Systems), AXL (386202; Biolegends), CD56 (362534), EPCAM (324222) or IgG control and then fixed in 2% PFA. Samples were analyzed on a BD LSRFortessa Flow Cytometer and data was analyzed using FlowJo 10.7.1.

### Knockdown of YAP1, RB1

YAP1-high LCNEC cell lines were transfected with either Validated Stealth Select RNAi or Smartpool RNAi for YAP1 or RB1 or Stealth RNAi Negative Universal Control at 10 nmol/L in OptiMEM using Lipofectamine according to the manufacturer’s protocol. Forty-eight hours after transfection, cells were harvested and lysates analyzed by western blot analysis for RB1 (SC-50; Santa Cruz Biotechnology), YAP1 (D8H1X; Cell Signaling), Vimentin (3932; Cell Signaling), Synaptophysin (SC-17750; Santa Cruz Biotechnology), and Actin (8457S; Cell Signaling).

### CRISPR/dCas9 activation system and transfection

For overexpressing YAP1, the cells were transfected with commercially available CRISPR/dCas9 lentiviral activation particles (Santa Cruz Biotechnology, Dallas, TX). Cells (2*10^5^) were cultured in the medium containing 10% FBS (antibiotic free) for 24h and infected with lentiviral vectors. After transduction, the cells were subjected to selection using puromycin (Thermo Fisher Scientific), blasticidin (InvivoGen, San Diego, CA), and hygromycin (Sigma-Aldrich, St. Louis, MO) to establish stable YAP1-overexpressing cell lines. YAP1 and PD-L1 (E13N; Cell Signaling), and Actin levels were verified by western blot analysis compared to the parental cell line.

### Drug treatment of cells

LCNEC cells were treated with DMSO, 1 μM cisplatin, verteporfin, MYF-01-37, XAV-939, or 5 μM decitabine for 72h. YAP1-high SCLC and LCNEC cell lines were treated with 1 μM trametinib for 72h. Cell lysates were harvested for western blot analysis. Antibodies used for western analysis include YAP1, AXL (8661S; Cell Signaling), Vimentin, vinculin (V9131, Sigma) as a loading control.

### Animal experiments

For patient derived xenograft models (PDX), fresh tumors were implanted subcutaneously into the flank of athymic nude mice. For cell line xenografts, one million cells in 1:1 PBS and Matrigel were injected subcutaneously into the flank of athymic nude mice. Treatment of the mice began when the tumors reached 120-150 mm^3^. Tumor volume and body weights were measured on all mice two times per week and calculated (width^2^ x length x 0.4). Vehicle or trametinib (100 mg/kg) were administered by oral gavage daily (five days per week). Vehicle or cisplatin (6 mg/kg) were administered intraperitoneally once per week. Maximum change in tumor growth from baseline was calculated for each model treated with cisplatin. All animals were maintained in accordance with the Institutional Animal Care and Use Committee of the M.D. Anderson Cancer Center and the NIH Guide for the Care and Use of Laboratory Animals.

### RPPA

Protein lysates from LCNEC cell lines, SCLC cell lines (H69, and H69CPR), and cell line xenograft tumors from H69 and H69 CPR were quantified and protein arrays were printed and stained, as described previously^22^. Images were quantified with MicroVigene 4.0 (VigeneTech, Carlisle, MA). The spot-level raw data were processed with the R package SuperCurve suite, which returns the estimated protein concentration (raw concentration) and a quality control score for each slide, as described previously^22^. Only slides with a quality control score of >0.8 were used for downstream analysis. The raw concentration data were normalized by median-centering each sample across all the proteins to correct loading bias.

### CAR T-cell co-culture experiments

LCNEC cell lines were transduced with a dual luciferase and GFP reporter, and GFP-positive cells were sorted and co-cultured with control non-transduced T cells (NT-T) and DLL3, CD56, or AXL CAR-Ts at an effector to target ratio (E:T) of 1:1 for 72h. Cytotoxicity was measured by bioluminescence. This system determines cytotoxicity by detecting cancer cell luminescence without separating CAR-Ts, which is otherwise difficult to perform in floating cell lines.

### Single-cell RNAseq Tissue Dissociation and Sequencing

Tissues were digested gently overnight at 37C using collagenase A (1 mg/ml). Red blood cell lysis was performed, as needed. Cells were counted and only samples with >50% viability were submitted for single-cell RNA sequencing at the MD Anderson’s CPRIT Single Core facility.

### Single-cell RNAseq Analyses

Raw data for single cell datasets were processed using the Cell Ranger v3.1.0^40^ to obtain the unique molecular identifier (UMI) data matrix. Cells with less than 300 detectable genes were filtered out. Samples from different batches were normalized and integrated following the sample integration pipeline in SEURAT v3^41,42^. The integrated datasets were processed using SEURAT, including selecting significant PCA components for dimension reduction, UMAP conversion and visualization, and density-based clustering for subpopulation discovery. The cell subpopulations were identified and annotated using SingleR package^43^ with manual curation. The gene expression levels were visualized on UMAP by either colored feature plot or binary plot. For each gene, the expression status is defined as positive if the cell has non-zero expression value, or negative otherwise. EMT score was calculated based on the EMT signature as described previously^25,44^.

### Statistical Analyses

All statistic and bioinformatics analyses were performed using R. Paired and un-paired two-sample t-tests were used for two group comparisons on paired and unpaired experimental design experiments. Pearson and Spearman correlations were used for correlating genomic and proteomic measurements, as well as correlating drug-screening data. In all the analyses, p<0.05 was considered statistically significant.

## Data Set Availability

Publicly available data were obtained from CCLE, DepMAP.

## Acknowledgements

We thank the patients who participated in this study, as well as their families. This work was supported by: The NIH/NCI CCSG P30-CA016672 (MD Anderson Flow Cytometry and Cellular Imaging Facility, the MD Anderson Bioinformatics Shared Resource, and the MD Anderson Institutional Tissue bank (ITB)); NIH/NCI T32-CA009666 (BZ); The University of Texas-Southwestern and MD Anderson Cancer Center Lung SPORE P50-CA070907 (JW, JVH, CMG, LAB); NIH/NCI R01-CA207295 (LAB); NIH/NCI U01-CA213273 (JVH, LAB); NIH/NCI R50-CA243698 (CAS); NIH/NCI U01-CA256780 (JVH, LAB); The Department of Defense LC210510 (LAB); the LUNGeveity Foundation 2020-02 (CMG); CPRIT RP210159 (CMG); NETRF (CMG); Through generous philanthropic contributions to The University of Texas MD Anderson Lung Cancer Moon Shot Program (JVH, LAB, CMG); and Rexanna’s Foundation for Fighting Lung Cancer (JVH, LAB, CMG). This study was supported by the Translational Molecular Pathology Immunohistochemistry and Digital Pathology Laboratory at the Department Translational Molecular Pathology, the University of Texas MD Anderson Cancer Center with special thanks to Wei Lu, Mei Jiang, Khaja Khan and Jianling Zhou for their technical assistance in the completion of this project. BBM is a TRIUMPH Fellow in the CPRIT Research Training Program (RP210028).

## Author Contributions

C.A.S. and C.M.G. conceived the project, analyzed and interpreted the data, and wrote the manuscript; C.A.S., R.W., B.Z., Y.Y. performed experiments and interpreted results; C.A.S., K.R., B.L.R., A.G.S. interpreted results; Y.X., L.D., L.S., B.B.M., P.D.R., L.S.S. and J.W. contributed to the analysis and interpretation of data; C.M.G., B.Z., M.S., V.Y.N., R.W., A.D., R.J.C., M.F., S.H.H., B.E.W., N.V., D.L.G., J.V.H., B.G., L.A.B. contributed to the acquisition of data, administrative, and/or material support. All authors contributed to the writing, review, and/or revision of the manuscript.

