## Supplemental figures for "YAP1 status defines two intrinsic subtypes of LCNEC with distinct molecular features and therapeutic vulnerabilities"

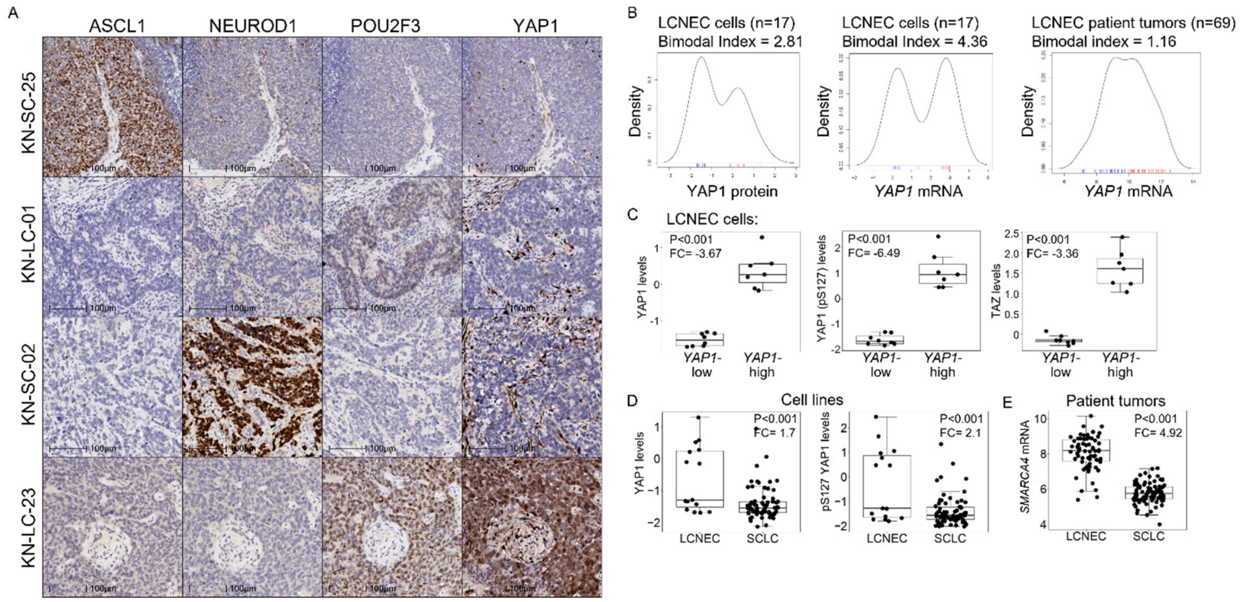


**Extended Data Figure 1. YAP1 is bimodal in LCNEC**. A, IHC for ASCL1, NEUROD1, POU2F3, and YAP1 in LCNEC patient tumors. B, YAP1 protein (left) and mRNA (middle) levels in cell lines and patient tumors (right) are bimodal. C, YAP1-high LCNEC cells have high levels of YAP1, phospho YAP1 (S127) and TAZ levels by RPPA. E, LCNEC cells have higher YAP1 and phospho YAP1 (S127) levels compared to SCLC. F, *SMARCA4* mRNA is higher in LCNEC tumors compared to SCLC, despite the frequent mutations. Scale bar = 100 μm.

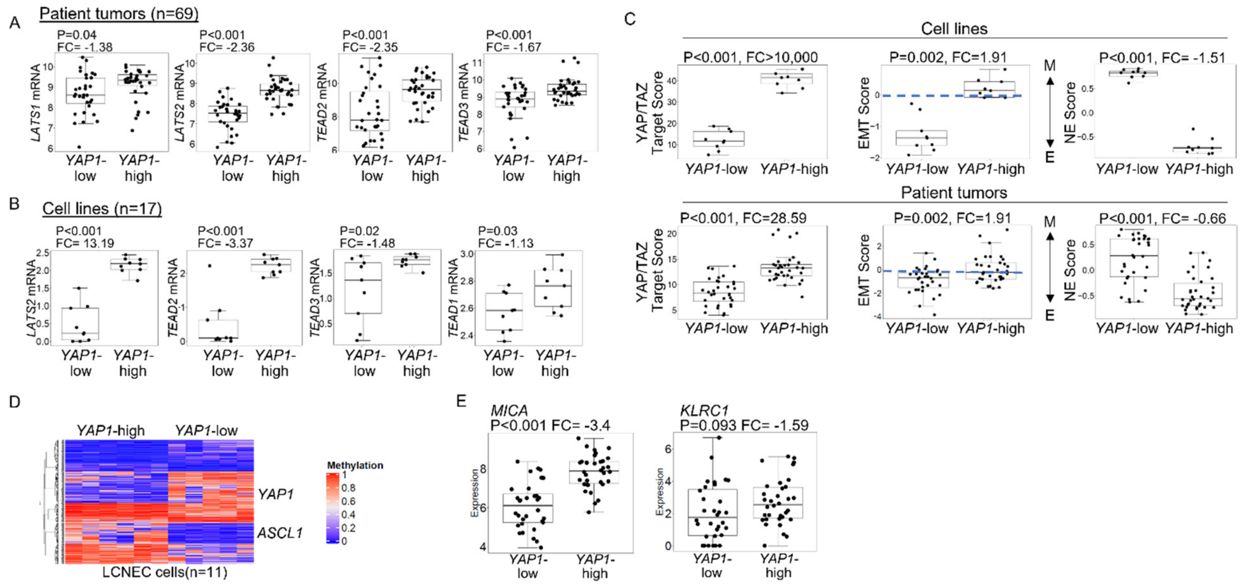


**Extended Data Figure 2. Unique features of YAP1-high LCNEC**. YAP1-high LCNEC patient tumors (A) and cell lines (B) express abundant hippo signaling pathway genes. C, YAP1-high cells (top) and tumors (bottom) have a higher YAP/TAZ target score (left) and EMT score (middle) and lower NE score (right). D, Heatmap of RRBS sequencing of YAP1-high and -low cell lines reveals *YAP1* and *ASCL1* are methylated in different subsets. E, Comparison of *MICA* and *KLRC1* (NKG2A) expression in YAP1-high and -low tumors.

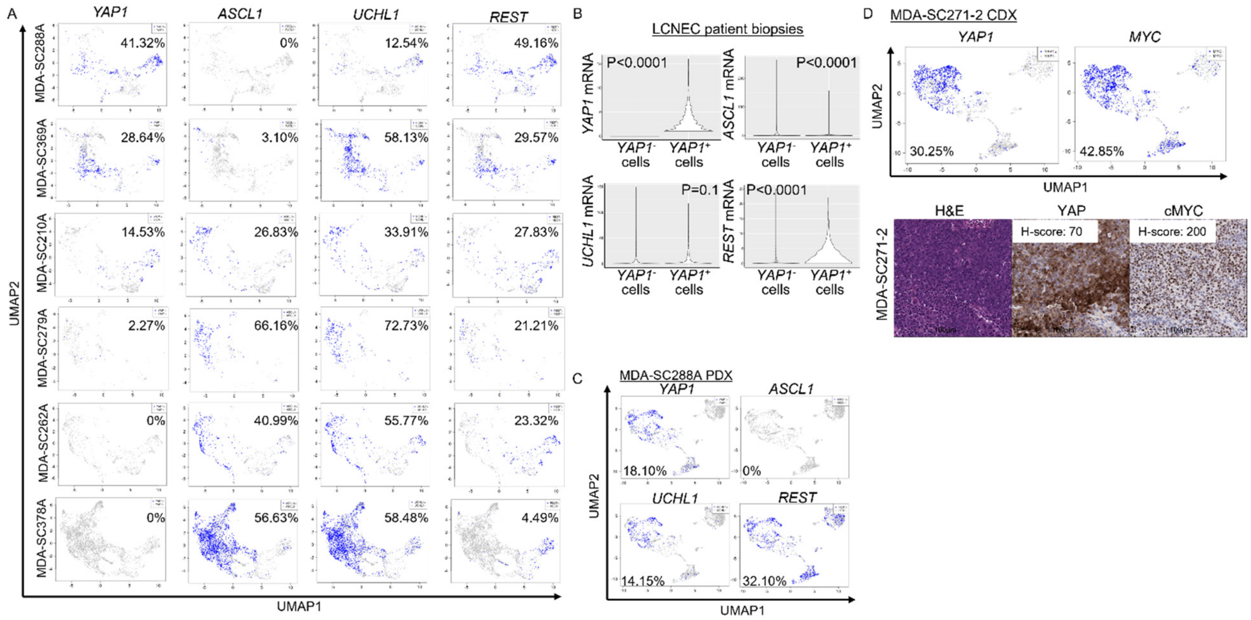


**Extended Data Figure 3. Single-cell transcriptional analyses of subtype-defining genes in LCNEC biopsies**. A, UMAP plots visualizing binary expression of *YAP1, ASCL1, UCHL1* and *REST* in 6 individual LCNEC patient cancer cell populations. B, Violin plots demonstrating *YAP1, ASCL1, UCHL1*, and *REST* expression in *YAP1*-positive and -negative cells. C, UMAP plots visualizing *YAP1, ASCL1, UCHL1*, and *REST* in MDA-SC288APDX. D, *YAP1* and *MYC* expression by single-cell RNAseq (upper) and corresponding H&E stained tumor with YAP1 and cMYC IHC (lower).

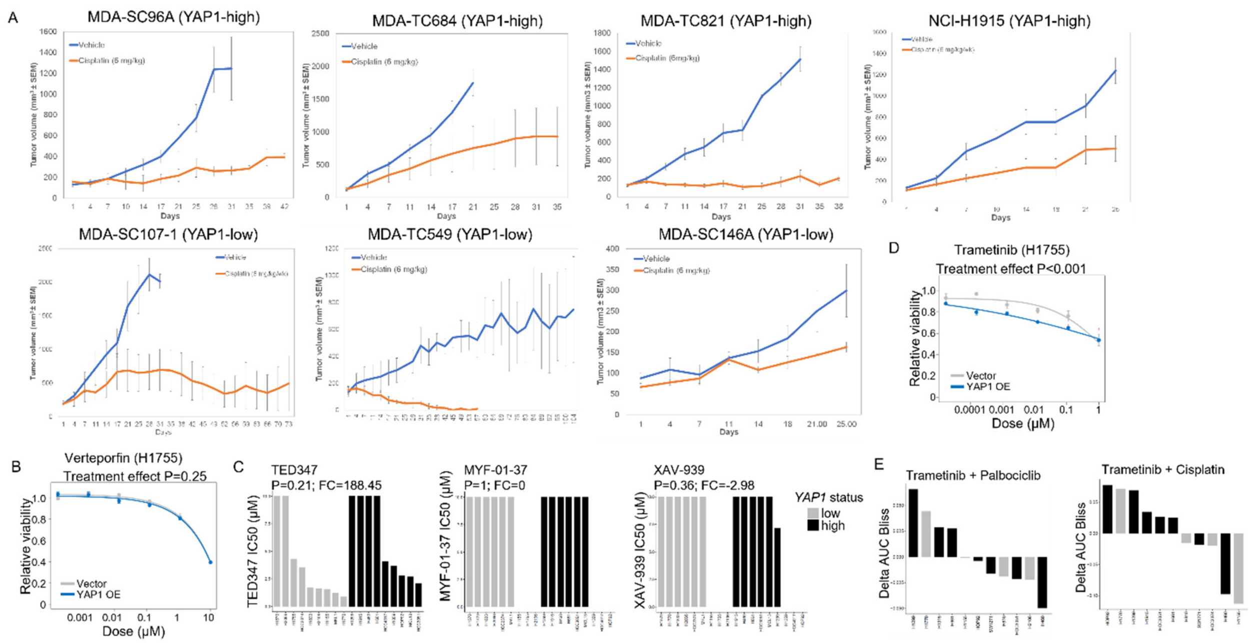


**Extended Data Figure 4. Drug response differences in YAP1-high and -low LCNEC preclinical models**. A, Tumor growth curves for YAP1-high and -low PDX models and cell line xenografts treated with 6 mg/kg cisplatin i.p. weekly. B, Overexpression of YAP1 did not change sensitivity to verteporfin. C, There is no difference in YAP1/TEAD inhibitor response based on YAP1 status. E, Combination of trametinib with either palbociclib or cisplatin did not improve response. F, Overexpression of YAP1 improved response to trametinib in a YAP1-low cell line.

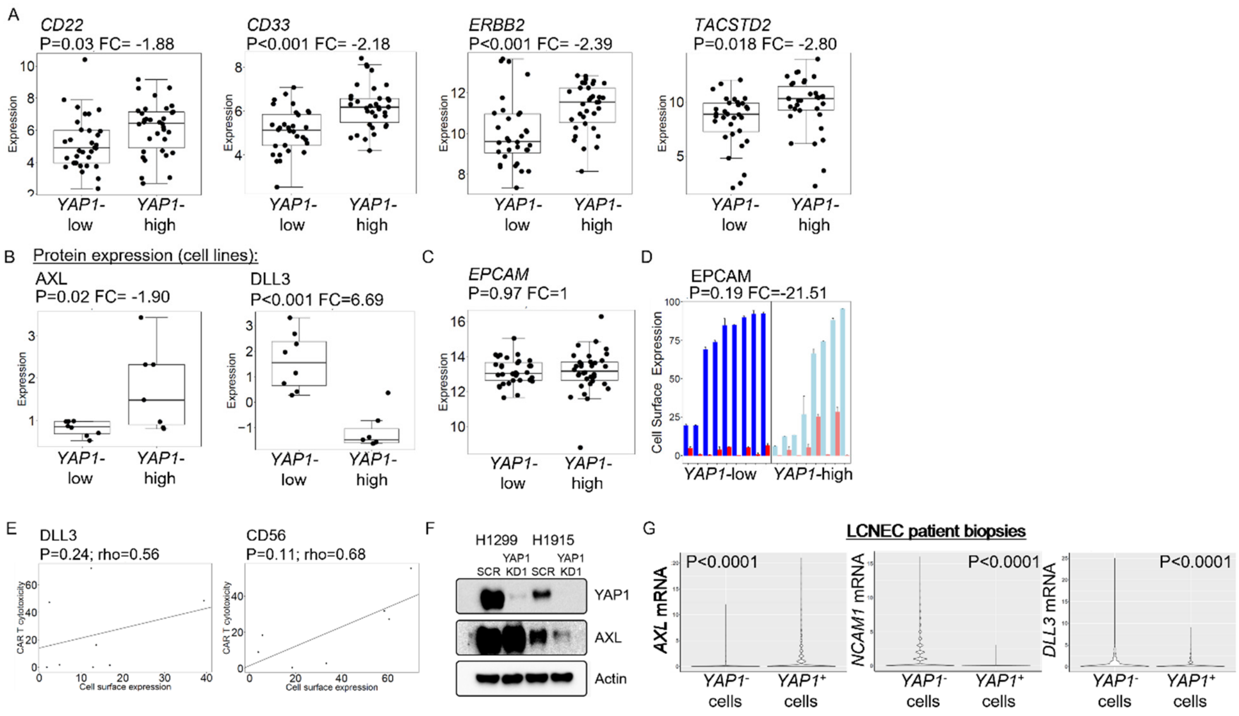


**Extended Data Figure 5. Surfaceome targets in LCNEC**. A, Expression of approved therapeutic surface targets in YAP1-high and -low LCNEC patient tumors. B, AXL protein is higher in YAP1-high and DLL3 is higher in YAP1-low LCNEC cell lines. C, *EPCAM* mRNA is higher in YAP1-high LCNEC patient tumors. D, Cell surface EPCAM is high in both YAP1-high and -low subsets of LCNEC. E, DLL3 and CD56 CAR-T cytotoxicity does not correlate with cell surface expression in LCNEC cell lines. F, Knockdown of YAP1 reduces AXL levels. G, Violin plots demonstrating AXL, NCAM1, and DLL3 in YAP1-positive and -negative cell populations.
